# Identification of novel natural product inhibitors of BRD4 using high throughput virtual screening and MD simulation

**DOI:** 10.1101/2022.11.25.517921

**Authors:** Soumen Barman, Snehasudha Subhadarsini Sahoo, Jyotirmayee Padhan, Babu Sudhamalla

## Abstract

Bromodomains are evolutionarily conserved structural motifs that recognize acetylated lysine residues on histone tails. They play a crucial role in shaping chromatin architecture and regulating gene expression in various biological processes. Mutations in bromodomains containing proteins leads to multiple human diseases, which makes them attractive target for therapeutic intervention. Extensive studies have been done on BRD4 as a target for several cancers, such as Acute Myeloid Leukemia (AML) and Burkitt Lymphoma. Several potential inhibitors have been identified against the BRD4 bromodomain. However, most of these inhibitors have drawbacks such as nonspecificity and toxicity, decreasing their appeal and necessitating the search for novel non-toxic inhibitors. This study aims to address this need by virtually screening natural compounds from the NPASS database against the Kac binding site of BRD4-BD1 using high throughput molecular docking followed by similarity clustering, pharmacokinetic screening, MD simulation, and MM-PBSA binding free energy calculations. Using this approach, we identified five natural product inhibitors having a similar or better binding affinity to the BRD4 bromodomain compared to JQ1 (previously reported inhibitor of BRD4). Further systematic analysis of these inhibitors resulted in the top three hits: NPC268484 (Palodesangren-B), NPC295021 (Candidine), and NPC313112 (Buxifoliadine-D). Collectively, our *in silico* results identified some promising natural products that have the potential to act as potent BRD4-BD1 inhibitors and can be considered for further validation through future *in vitro* and *in vivo* studies.

## Introduction

DNA accessibility and transcriptional activation in the eukaryotic genome are often regulated by reversible posttranslational modifications (PTMs) such as acetylation, phosphorylation, methylation, and ubiquitylation on histone proteins.^*1*^ The combinatorial readout of these PTMs are crucial in modulating the gene expression without altering the DNA sequence.^*2*^ There are three classes of epigenetic proteins: writers, readers, and erasers, whose coordinated activities are responsible for different patterns of PTMs on the histone tail eventually regulating gene expression.^*3*^ Since the advent of epigenetics as a discipline, one of the most studied PTMs has been histone acetylation, which corresponds to the addition of an acetyl group to the ε-amino side chain of lysine on the N-terminal tails of histones.^*4*^ This mark is recognized by members of the bromodomain family which are evolutionarily conserved protein interacting modules found in 46 diverse human proteins, containing a total of 61 bromodomains.^*5*^ Although all bromodomain proteins have significant sequence differences, they all share high structural similarities and same folding patterns.^*5*^ A typical bromodomain motif consists of four left-handed alpha helices (αZ, αA, αB, and αC), connected by ZA and BC loops which is responsible for substrate specificity (Figure 1).^*6*^ Recognition of acetylation mark on histone tails by bromodomains is essential in regulating chromatin structure, replication, transcription, and cell cycle progression.^*7*, *8*^

**Figure 1.**
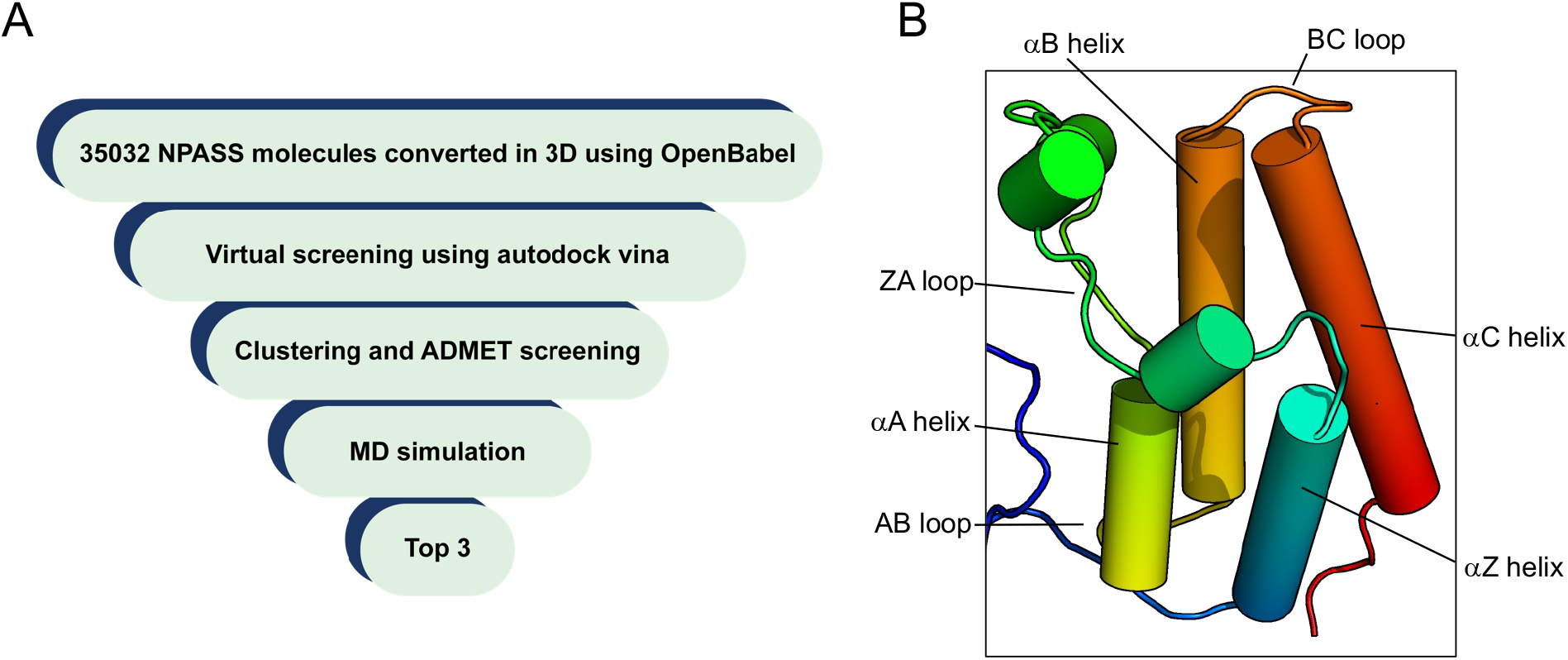
(A) Flowchart summarizing the systematic computational strategy to select BRD4-BD1 inhibitors from a library of NPASS (Natural Product Activity and Species Source) data base. (B) The archetypical four-helix bundle topology of the bromodomain.

Based on domain organization, bromodomains can be classified into the BET (Bromodomain and extra-terminal domain) and non-BET family members. The BET family of bromodomain proteins consists of four members, BRD2, BRD3, BRD4, and BRDT, that contain two N-terminal bromodomains and an extra-terminal domain in the C-terminus.^*9*^ Among them BRD4 is the most studied bromodomain and has gained wide attention recently, as it is highly misregulated in various diseases, including cancers.^*10*^ In the context of breast cancer, BRD4 helps in the epithelial to mesenchymal transition by regulating the expression of various EMT transcription factors.^*11*^ BRD4 binds to acetylated TWIST (an EMT TF) and forms a complex on the promoter and enhancer of metastasis-promoting genes in basal-like breast cancer.^*12*^ In addition, it is also associated with many other types of malignancies, such as melanoma, glioblastoma, breast cancer, and prostate cancer, and non-small cell lung cancer, coupled with other diseases including inflammation and HIV.^*13*, *14*^

In recent years, attempts have been made to develop small-molecule epigenetic target modulators for cancer therapeutics through combination of *in vitro* and in-silico methods.^*15*^ Filippakopoulos et al. have investigated the selective inhibition of BET using small-molecule based ligands and developed a highly effective methyltriazolodiazepine-based inhibitor JQ1, which has a high affinity for the bromodomains of BRD4 (IC50 value of 77 nM for BD1 and 33 nM for BD2).^*16*^ Another small molecule-based ligand, I-BET762, was screened and developed by GlaxoSmithKline (GSK), and the IC50 of this compound varies from 32.5 to 42.5 nM for the BET BRDs.^*17*^ A 3,5-dimethylisoxazole-based derivative, named I-BET151, also exhibits a binding affinity of 200-790 nM for the BET-BRDs.^*18*^ Emily J. et al. developed ABBV-075, a potent, selective BET inhibitor, which targets BRD2, BRD4, and BRDT and its phase I clinical trials have been performed recently to evaluate its safety and pharmacokinetics in advanced solid and hematological tumors.^*19*^ The role of JQ1, I-BET762, and I-BET151 as bromodomain inhibitor have been investigated in cancers and inflammation.^*20*, *21*^ Recently Conway et al. reported that I-BET762 is currently under clinical trial to treat NUT-midline carcinoma (NMC).^*22*^

While the results of these BRD4 inhibitors have been quite promising in preclinical trials, several issues have been identified with these drug candidates. Among them the most promising inhibitor JQ1, could target conserved bromodomain binding pocket residues, implying that the inhibitors are not specific to BRD4 and might act on other members of the bromodomain family, resulting in significant unwanted side effects.^*23*^ Secondly, JQ1 has been recently reported to be toxic and trigger cell death through the intrinsic apoptotic pathway in neuronal derivative (ND) cells.^*24*^ This greatly warrants the need to identify fully potent and specific drug molecules against BRD4 that are safe and do not cause damage to non-target cells and tissues.

In the last few decades, high throughput virtual screening (HTVS) of drugs has been steadily replacing traditional methods of drug development with more preferred early-stage drug screening.^*25*^ Molecular docking and molecular dynamic simulation approaches have been used to interpret binding mode of inhibitors with receptor proteins in order to design more selective drugs.^*26*, *27*^ Structure-based HTVS, along with molecular docking and molecular dynamic simulation approaches, can help screen large libraries of ligand molecules and narrow them down to a few high potential drug candidates, which can be further validated *in vitro* and *in vivo*. In this study, we have adopted the approach of HTVS to find potential natural product inhibitors against BRD4-BD1. Natural products (NPs) are secondary metabolites derived from natural sources, usually safer substitutes of synthetic drug molecules.^*28*, *29*^

In this study, we screened 35,032 NPs from the natural product activity and species source (NPASSv1.0) database against BRD4-BD1.^*30*^ Molecular docking via AutoDock Vina was initially employed to narrow down the extensive library of NPs. The shortlisted candidates were clustered and screened based on their structural similarity and pharmacokinetic properties using ChemMine tools and the SwissADME server, respectively.^*31*–*33*^ MD simulation was conducted for the top five drug-like molecules, followed by MM/PBSA energy calculations to find the top three NPs as inhibitors.^*34*^ The MD simulation and the calculated binding energies of these potent inhibitors were also compared with that of JQ1.

## Materials and Methods

A schematic representation of the workflow adopted for the current study has been provided in Figure 1.

### Protein preparation

The crystal structure of the BRD4-BD1 protein complex with inhibitor JQ1 (PDB: 3MXF)^*16*^ was obtained from the protein data bank (RCSB). The structure was then minimized using the minimize structure module (1000 steps of steepest descent) using AMBER ff14SB force field in UCSF Chimera and Gasteiger charges were added.^*35*^ Next, the crystal structure of the protein target was prepared for docking by removing water molecules, ligands, and ions; missing residue or atoms were added. Finally, polar hydrogens and Kollmann-united charges were added to the protein using MGL AutoDock Tools (ADT), and the prepared protein was exported as a PDBQT for docking studies.

### Preparation of ligand library

A ligand library of all 35032 compounds in the Natural Product Activity and Species Source (NPASS) database (available at http://bidd2.nus.edu.sg/NPASS) was prepared. Two-dimensional structures of each compound were downloaded in structure-data file (SDF) format and converted into 3D ligand structure using the –gen3d module of Open Babel.^*36*^ Open Babel was further used to separate individual ligand files, add hydrogens, assign rotatable hydrogen bonds, and Gasteiger-Marsili charges to each of the ligands to prepare the ligand files. These ligand files were finally saved in PDBQT format to be used for subsequent molecular docking. The JQ1 inhibitor from the BRD4-BD1-JQ1 complex (PDB file: 3MXF) was also prepared by following the similar procedure described above and this structure was used as a control for docking studies.

### Molecular docking and virtual screening

The ligand binding site of the BRD4-BD1 receptor was identified by analyzing the crystal structure of the BRD4-BD1-JQ1 complex. Next, the ADT (AutoDock tools) was used to generate a grid box. The grid box dimensions were set to 42 × 42 × 45Å along the *X, Y*, and *Z* axes with a default grid spacing of 0.375 Å, which covered the entire protein. Molecular docking was performed using AutoDock Vina 1.1.2. To validate the docking method and the accuracy of the selected grid box dimensions, we initially performed docking studies with the reported inhibitor JQ1. We docked the JQ1 into the binding pocket of the BRD4-BD1 receptor. Once validity was confirmed, ten best-ranked docked conformations were generated for each compound from the ligand library with the protein. The docking scores of these poses were further verified with the SwissDock tool to reduce false positive hits. The ligand poses with docking scores less than JQ1’s docking score was chosen for further screening. The potential inhibitors were sorted according to their docking scores. Molecular docking was done using written in-house scripts. All visualizations of crystal structures were performed using the PyMOL software package.

### Ligand similarity

ChemMine tool was used to analyze the chemical and structural similarity between ligand compounds.^*32*^ Tanimoto coefficient (Tc) was employed to calculate the chemical similarity between the ligand pairs.^*37*^ Given two compounds, X and Y, Tanimoto (X, Y) = z/(x+y-z), where x represents the number of bits set to 1 in X, y represents the number of bits set to 1 in Y, and z represents the number of bits set to 1 in both. Thus, a similarity matrix is obtained with a range from 0 to 1, with a higher value indicating more similarity between the ligand pair and vice versa. Ligands were also classified based on their structure. OpenBabel 2.4.0 was employed to create SMILES structures from the SDF files of ligand compounds which was then uploaded into the ChemMine tool and multidimensionally scaled to get different clusters of structurally similar ligand compounds.

### Pharmacokinetic screening

Pharmacokinetic properties assist in the early stages of drug discovery by evaluating identifying the safety and effectiveness of the compounds. Thus, pharmacokinetic screening of top compounds was conducted with the SwissADME server.^*33*^ Compounds showing the least violations in their pharmacokinetic properties are considered good ligands and promising drug candidates. Hence, the top 5 ligand compounds from this screening were selected for further analysis.

### Molecular dynamics simulation

Computational modeling and MD simulation were performed by following the procedure as previously described with slight modifications.^*38*^ The top 5 best-ranked docked conformations of BRD4-BD1 and ligand complexes were subjected to 100 ns of MD simulations with GROMACS (version 2018.3) software package using the CHARMM27 force field.^*39*, *40*^ In order to set up MD simulations for the protein-ligand complexes, the topology parameters of the BRD4-BD1 and natural product ligands were created using GROMACS and CGenFF server, respectively.^*41*^ The prepared protein-ligand complexes were then solvated in a dodecahedron box with a distance of 1.0 nm between the complex and the edge of the solvated box. The solvated system was neutralized by adding chloride ions in the simulation. The system was energy minimized using the LINCS constraints and steepest descent algorithms followed by system equilibration under NVT and NPT ensembles to ensure the complex’s steric clashes or geometry. Final MD simulation of bromodomain and inhibitor complexes were carried out at 300 K temperature, 1 atm pressure, and 2 fs time steps for 100 ns. The final MD trajectories were analyzed to calculate the RMSD (root mean square deviation), RMSF (root mean square fluctuation), Radius of gyration (Rg), number of hydrogen bonds and distance measurements using the standard GROMACS functions. The MD simulations and results analysis were performed on the DIRAC supercomputing facility of IISER Kolkata.

### MM-PBSA binding free energy calculations

MM-PBSA (Molecular Mechanics Poisson Boltzmann Surface Area) binding free energy calculations were performed as previously described.^*42*^ The g_mmpbsa tool was used to calculate the free energy of binding of protein-ligand complexes from MD trajectories.^*43*, *44*^ Post simulation, MM-PBSA calculations of the free energy of binding have been used extensively to screen inhibitors and found to correlate reasonably well with experimental results.^*34*, *45*, *46*^ This module estimates the Gibbs free energy of binding using the MM-PBSA method as described by the following equations:

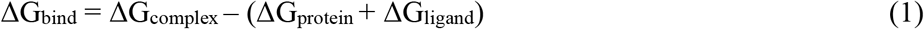

Where, ΔG_complex_, ΔG_protein_ and ΔG_ligand_ represent the total free energy of the protein-ligand complex and the free energies of the isolated protein and ligand in the solvent, respectively.

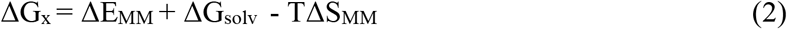

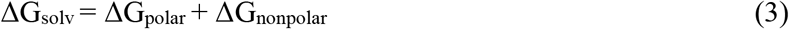

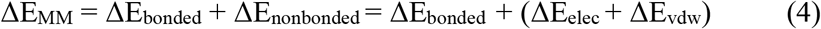

ΔG_x_ is the free energy, and TΔS refers to the entropic contribution to the free energy in the vacuum, where ΔT and ΔS describe the temperature and entropy, respectively. ΔG_solv_ is the free energy of solvation, where ΔG_polar_ and ΔG_nonpolar_ are the electrostatic and non-electrostatic contributions to the solvation-free energy. Vacuum potential energy, ΔE_MM_, includes the energy of both bonded and non-bonded interactions. The g_mmpbsa also allows the decomposition of the total binding free energy into contributions made by individual residues, giving us insight into potential residues driving the recognition and binding process.

### Toxicity analysis of top natural product inhibitors

The relative toxicity of top drug candidates was analyzed via the ProToxII server.^*47*^ This popular web server efficiently predicts various toxicity endpoints by integrating molecular similarity, fragment tendency, and fragment similarity methods. The server also predicted oral toxicity based on two-dimensional (2D) similarity analysis to compounds with a known mean lethal dose (LD50). The set used for the prediction consists of approximately 38,000 unique compounds with known oral LD50 values measured in rodents.

## Results

### Validation of molecular docking protocol using JQ1 as the reference inhibitor compound

Molecular docking can be used to model interactions between proteins and small molecules at the atomic level, allowing us to characterize the behaviour of small molecules in the binding site of the target proteins and elucidate fundamental biochemical processes.^*48*^ The docking process hinges on two basic steps: predicting the conformation, position, and orientation of a ligand in the protein binding site (altogether termed the ligand pose) and calculating the binding affinity of the provided ligand for the protein. This provides a nascent understanding of the potential ligand binding partners and can be used as a basis to screen and predict potential inhibitors of a targeted protein.

JQ1, a well-studied inhibitor of BRD4, was taken as the reference for our study. The validity of the molecular docking protocol utilized for this study was tested by redocking JQ1 with BRD4-BD1 in AutoDock Vina and the resultant docked structure compared with the documented co-crystal structure of BRD4-BD1-JQ1 complex (PDB ID: 3MXF). Both the docked and co-crystal structures of BRD4-BD1 and JQ1 were found to interact with the critical acetyllysine binding pocket residue N140 (Figure 2). Superimposition in PyMOL revealed a root mean square deviation of 3Å between the docked and the crystal structures of BRD4-BD1-JQ1 complexes confirming that the docking protocol is suitable for screening novel inhibitors against the BRD4-BD1.

**Figure 2.**
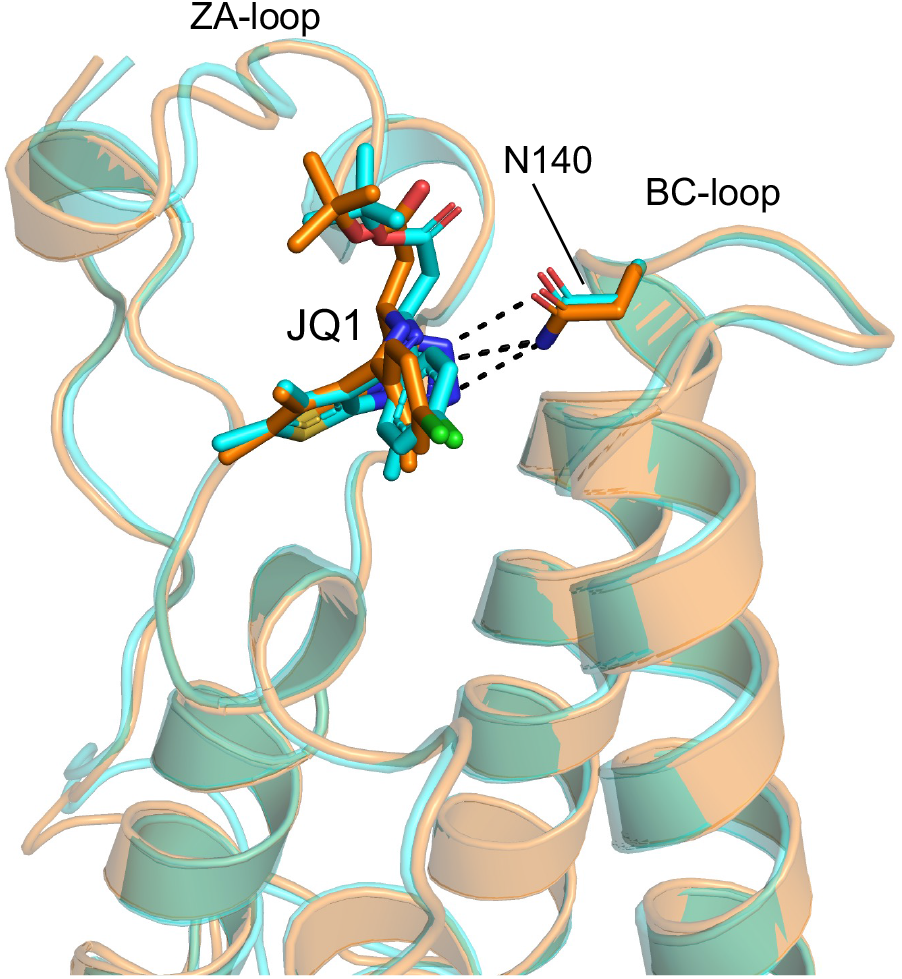
Structural superimposition of best ranked docked conformation of BRD4-BD1-JQ1 (cyan color) on the crystal structure of JQ1 inhibitor in complex with the BRD4-BD1 (PBD ID: 3MXF) (orange color) shows that the inhibitor binding pose is very much similar to the crystal structure and forms a critical hydrogen bond with the N140 residue of BRD4 bromodomain.

### Some natural products exhibit better docking scores compared to the well-known BRD4 inhibitor, JQ1

The main goal of the present study is to develop selective and potent natural product inhibitors against the BRD4-BD1. BRD4 is the most studied bromodomain-containing protein, and JQ1 is reported as a potential inhibitor of BRD4-BD1. In the initial stage of the docking, 2D coordinates of the compounds were downloaded from the NPASS server and converted into 3D structures using Open Babel. Then the 3D coordinate structures are further converted into pdbqt for docking using AutoDock Vina.

The natural product library containing 35032 compounds was screened against the BRD4-BD1, and the obtained docking scores were compared with the docking score of BRD4-BD1 and JQ1 inhibitor. Docking of JQ1 inhibitor with the BRD4-BD1 had yielded a score of −7.9 kJ/mol in AutoDock Vina and hence, inhibitors having a docking score lower than JQ1 (−7.9 kJ/mol) were identified and listed separately (Table 1 and Figure S1). The top 10 compounds were selected, and the overall docking score for these NPs ranged from −10.3 kJ/mol to −11.5 kJ/mol. The docking scores of NPC183 (2-[4-[5-(5,7-Dihydroxy-4-Oxochromen-2-Yl)-2-Methoxyphenoxy] Phenyl]-5,7-Dihydroxychromen-4-One), NPC12461 (Ugonin L), NPC28280 (Cyclopamine), NPC103230 (Decaturin D), NPC268484 (Palodesangren B), NPC290830 (Sotetsuflavone), NPC295021 (Candidine), NPC303485 (Podocarpusflavone A), NPC313112 (Buxifoliadine-D) and NPC473313 (4,7,8-Trihydroxy-2,3,3-Trimethyl-11-(3-Methylbut-2-Enyl)-2H-Furo[3,2-B] Xanthen-5-One) were found to be −10.8 kJ/mol, −10.4 kJ/mol, −10.3 kJ/mol, −10.4 kJ/mol, −11 kJ/mol, −10.3 kJ/mol, −11.5 kJ/mol, −10.4 kJ/mol, −10.6 kJ/mol and −10.3 kJ/mol, respectively (Table 1). NPC295021 has the best docking score of −11.5 kJ/mol among all the above listed compounds.

**Table 1.** Summary of the top 10 scored NPASS compounds against BRD4-BD1 including JQ1

| Molecule Common Name | NPASS | Pubchem ID | No. of H-Bonds | Interacting Residues | No. of non-covalent interactions | Interacting amino acids | Vina Score | Swiss Dock score |
| --- | --- | --- | --- | --- | --- | --- | --- | --- |
| **JQ1** |  | 46907787 | 1 | N140 | 7 | W81, P82, F83, L92, L94, Y139, I146 | -7.9 | -8.46 |
| **2-[4-[5-(5,7-Dihydroxy-4-Oxochromen-2-Yl)-2-Methoxyphenoxy]Phenyl]-5,7-Dihydroxychromen-4-One** | NPC183 | 5384799 | 5 | P82, D88, Y97, M132, N140 | 6 | P82, D88, L92, L94, Y97, I146 | -10.8 | -9.69 |
| **Ugonin L** | NPC12461 | 10365741 | 2 | M105, N135 | 6 | P82, F83, V87, L92, N135, I146 | -10.4 | -8.79 |
| **Cyclopamine** | NPC28280 | 442972 | 1 | M132 | 8 | P82, F83, V87, L92, L94, Y97, Y139, I146 | -10.3 | -8.57 |
| **Decaturin D** | NPC103230 | 11178957 | 2 | Y97, K141 | 5 | F83, V87, L94, Y139, I146 | -10.4 | -8.48 |
| **Palodesangren B** | NPC268484 | 10648988 | 3 | Y97, N140, I146 | 7 | W81, P82, F83, V87, L92, Y97, I146 | -11 | -8.79 |
| **Sotetsuflavone** | NPC290830 | 5494868 | 4 | Y97, N140, D145, I146 | 4 | F83, L94, Y139, I146 | -10.3 | -9.24 |
| **Candidine** | NPC295021 | 5320815 | 2 | P82, N140 | 4 | P82, F83, L92, I146 | -11.5 | -7.48 |
| **Podocarpusflavone A** | NPC303485 | 5320644 | 3 | Y97, N140, D145 | 4 | F83, L94, Y139, I146 | -10.4 | -9.31 |
| **Buxifoliadine-D** | NPC313112 | 10428666 | 3 | P82, Y97, N140 | 4 | P82, F83, Q85, L92 | -10.6 | -9.03 |
| **4,7,8-Trihydroxy-2,3,3-Trimethyl-11-(3-Methylbut-2-Enyl)-2H-Furo[3,2-B]Xanthen-5-One** | NPC473313 | 25028557 | 2 | Y97, N140 | 7 | P82, F83, Q85, L92, Y139, I146 | -10.3 | -7.75 |

The top-scored docking conformations of 3D and 2D interaction analysis revealed the formation of hydrogen bonding and hydrophobic interactions between the natural product compounds and the critical binding pocket residues of BRD4-BD1 (Figure 3). The most common interactions between NPs and BRD4-BD1 constitute hydrogen bonding with the N140 and Y97 and hydrophobic interactions with W81, P82, F83, L92, L94, Y139, and I146 (Table 1 and Figure 3). Furthermore, to compare and validate the docking scores of these potential inhibitors, we performed additional docking calculation using the SwissDock server. It was found that the docking scores obtained through the SwissDock server are very similar to the vina docking scores. (Table 1).

**Figure 3.**
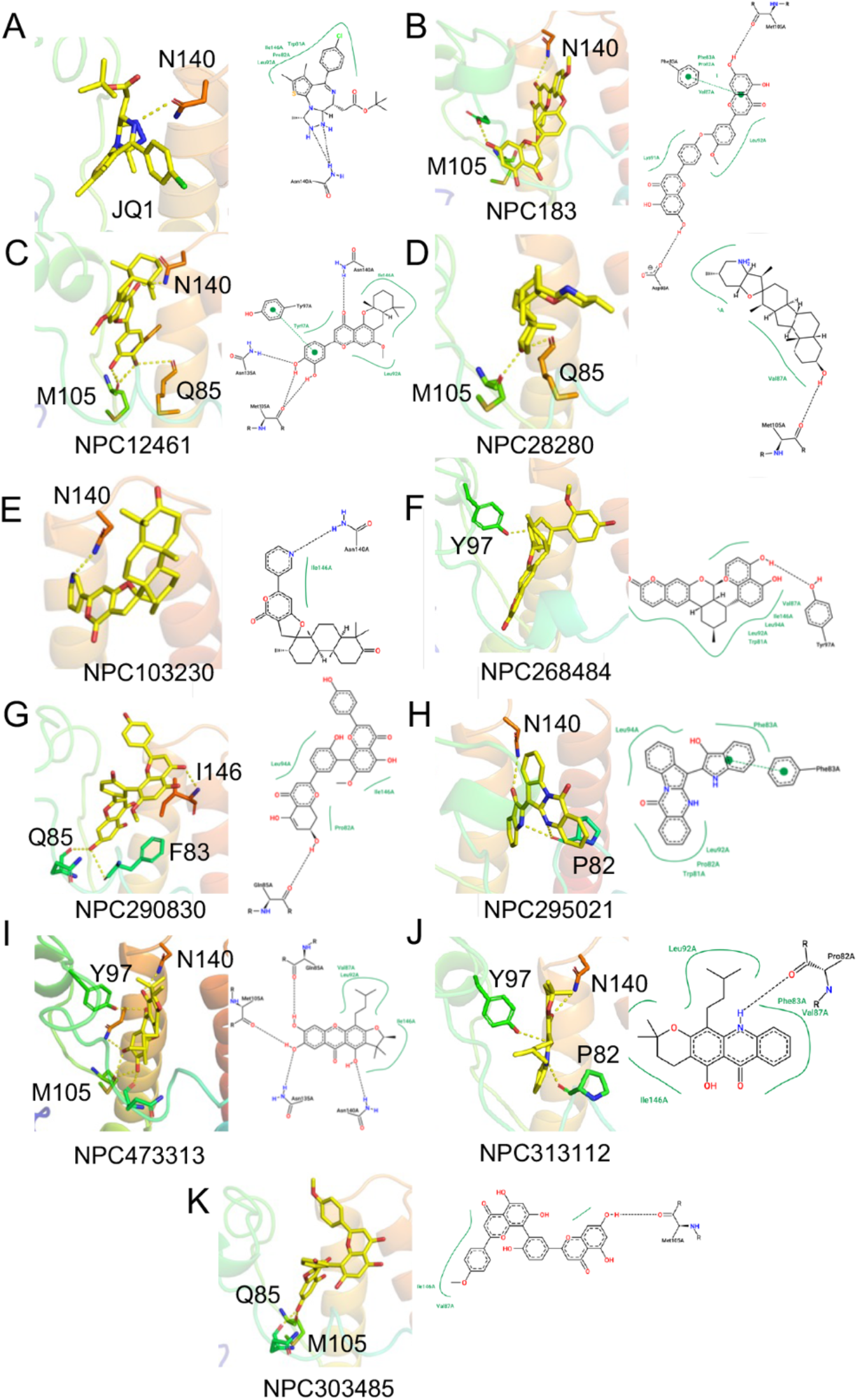
The best ranked natural product inhibitors (top 10 NPs) in complex with the BRD4-BD1. The interaction between the bromodomain binding pocket residues and ligands are highlighted.

### Similarity analysis and physicochemical properties

Even though the newly identified natural products exhibit favourable docking scores and hydrogen bonding interactions with BRD4-BD1, they need to be unique and measure up to drug likeliness standards to be considered critical drug leads. Clustering and analyzing the top 10 NPs in ChemMine tools yielded a multidimensional scaling (MDS) scatter plot based on structural similarity using the Tanimoto coefficient with a cutoff of 0.4 (Figure S2). A total of two groups with similarity coefficient <0.4: the first group with three compounds named NPC183, NPC290830 and NPC303485 and a second group with two compounds, NPC313112 and NPC473313 were depicted, resulting in a total of seven structurally distinct chemical compounds within the top 10 NPs (Figure S2). The hierarchical clustering of joe lib descriptor heatmap shows the comparative structural feature between these compounds (Figure S3).

Furthermore, predictions of the physicochemical properties of the above selected compounds are essential for evaluating their drug-likeness and lead likeness *in silico*, making them a critical component of drug lead studies. For example, the ‘Lipinski Rule of Five’ filters compounds with ≤ 5 hydrogen bond donors, ≤ 10 hydrogen acceptors, a molecular weight ≤ 500 daltons, an octanol-water partition coefficient log P≤5 and, molar refractivity between 40-130. The total polar surface area (TPSA) is another drug likeness property, which needs to be less than 140 Å to represent good absorption capacity of the drug lead.

Analysis of the calculated physicochemical properties through the SwissADME online tool revealed that all the compounds have a log p-value of less than 5, suggesting that these compounds are likely to be absorbed effectively. NPC183 has a molecular weight of more than 500 daltons and a total polar surface area (TPSA) of 170 Å (Table 2). Another topological parameter checked is the number of rotatable bonds. All compounds have a minimum of 1 rotatable bond, except NPC28280, with no rotatable bonds. Based on these analyses, we found that NPC183 and NPC28280 violate the Lipinski rule. The other 5 NPs; NPC12461, NPC103230, NPC268484, NPC295021, and NPC313112, have drug-likeliness properties with minimum violations and thus, check the pre-requisites to be potential drug candidates (Table 2). These top five compounds were further subjected to MDS for further evaluation.

**Table 2.**
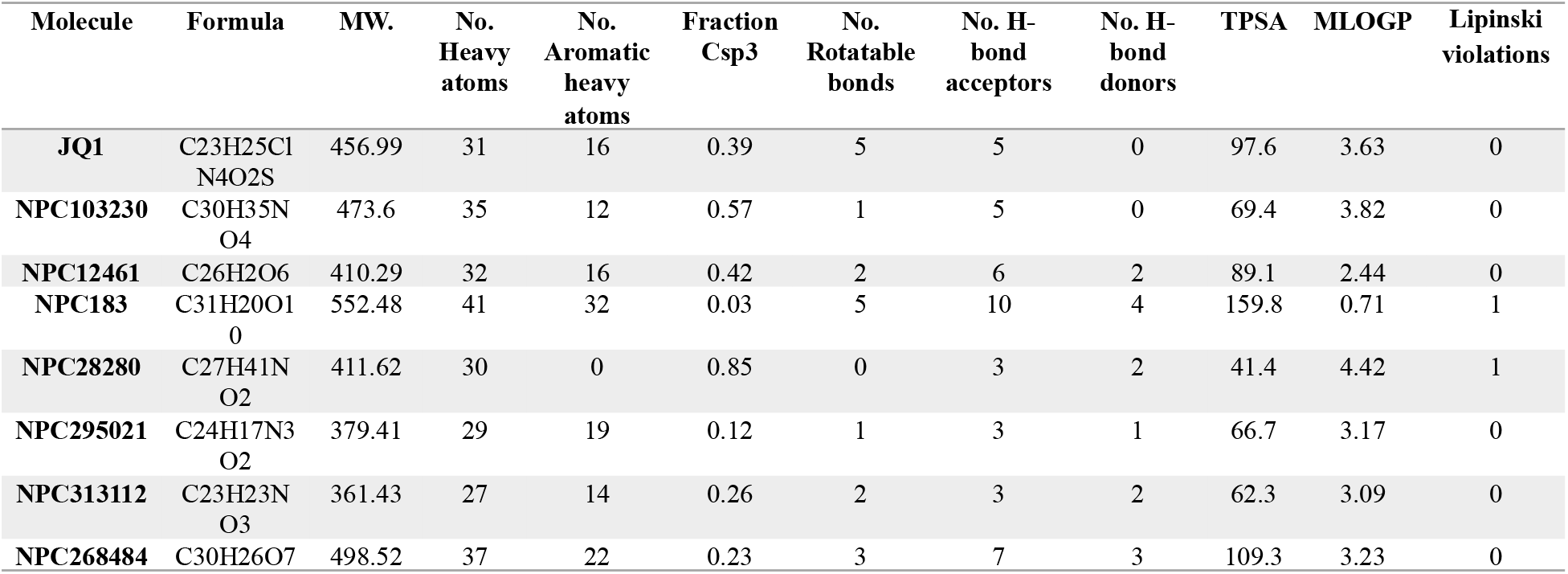
Drug-likeness property analysis of top 7 phenolic compounds from NPASS

### Molecular dynamics simulation analysis

Next, we subjected six complexes (BRD4-BD1-NPs) for molecular dynamic simulation, including JQ1 as the reference molecule. The MD simulation aimed to get a more precise and realistic idea of the interactions between the protein and ligand complexes and their binding mode. In this study, the RMSD (Root mean square deviation), RMSF (Root means square fluctuations) of the Cα backbone and the number of hydrogen bonds between the protein and drug molecule was used to represent the stability and dynamics of the complex under normal physiological conditions over 100 ns (Figure 4). The lower the RMSD and RMSF values, the better the binding, and vice versa. Radius of gyration (Rg) is calculated to understand the compactness, stability and folding of structure. The strength of inhibitors binding with BRD4-BD1 was studied using pair distance analysis and the number of hydrogen bonds between the protein and drug candidate in the complex is directly proportional to its stability.

**Figure 4.**
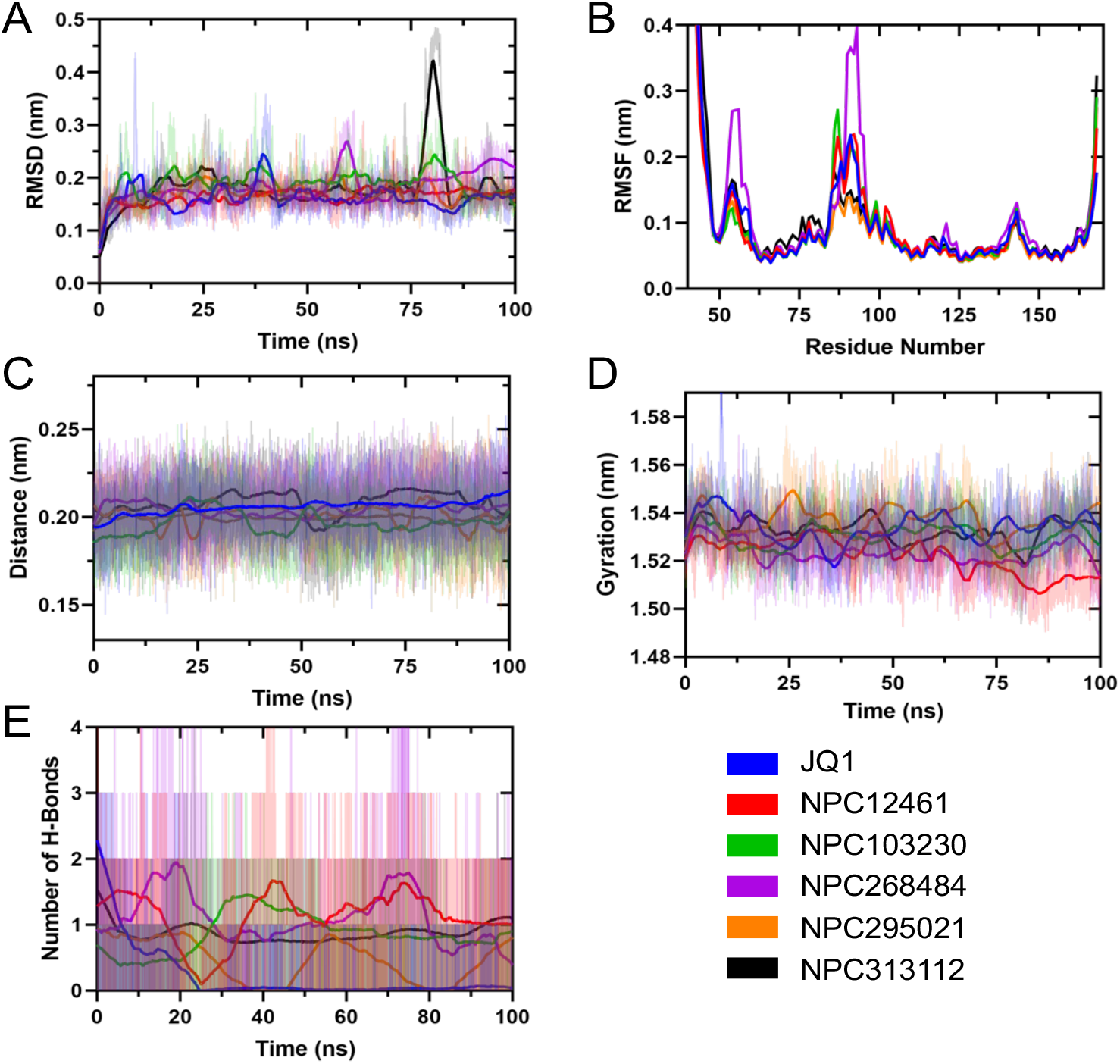
(A-E) Root-mean-square deviation (RMSD), root-mean-square fluctuation (RMSF), paired distance, radius of gyration (Rg), and hydrogen bonding of protein-ligand complexes were plotted over 100 ns of MD simulation.

The average of the backbone RMSD, RMSF, distance, Rg, and the number of hydrogen bonds in different complexes, is represented in Figure 4A-E. RMSD analysis revealed that the compound NPC313112 shows the highest, NPC103230 and NPC268484 show moderate, and NPC12461 and NPC295021 show similar fluctuations when compared to JQ1, respectively (Figure 4A and B). RMSF analysis reveals that the NPC268484 and NPC313112 complex show the most fluctuation throughout the simulation, while the other three are similar to JQ1 (Table 3). The distance between BRD4-BD1 protein and natural product inhibitors was extracted from the MD trajectory from the whole simulation and plotted against time (Figure 4C). The global minimum distance between protein and NPs is ~0.2 nm which explains that the number of hydrogen bonds formed between them is significant enough to maintain a constant minimum distance of ~2 Å (Table 3). All the complexes have a steady average Rg of ~1.5 nm over the 100 ns MD simulation (Figure 4D). The average number of H-bonds formed between BRD4-BD1 and different NPs varies from 0.5 to 1.2, which is higher than the H-bonds formed by JQ1 in the MDS (Figure 4E and Table 3).

**Table 3.** The mean ± SD of MD parameters of BRD4-BD1 and NPs interaction

|  | **JQ1** | **NPC12461** | **NPC103230** | **NPC268484** | **NPC295021** | **NPC313112** |
| --- | --- | --- | --- | --- | --- | --- |
| RMSD (nm) | 0.1625 ±<br>0.0212 | 0.1653 ±<br>0.01408 | 0.1902 ±<br>0.02275 | 0.1821 ±<br>0.02933 | 0.1705 ±<br>0.01646 | 0.1808 ±<br>0.04664 |
| RMSF (nm) | 0.09221 ±<br>0.07305 | 0.09192 ±<br>0.06828 | 0.09264 ±<br>0.07467 | 0.1111 ±<br>0.09524 | 0.08202 ±<br>0.06235 | 0.0997 ±<br>0.08839 |
| Rg (nm) | 1.534 ±<br>0.006055 | 1.522 ±<br>0.007312 | 1.53 ±<br>0.004373 | 1.523 ±<br>0.004321 | 1.537 ±<br>0.00541 | 1.533 ±<br>0.004585 |
| H-Bond | 0.2156 ±<br>0.423 | 1.13 ±<br>0.371 | 0.8632 ±<br>0.298 | 1.104 ±<br>0.3816 | 0.4605 ±<br>0.319 | 0.8713 ±<br>0.1331 |
| Paired<br>Distance<br>(nm) | 0.2057 ±<br>0.01193 | 0.1966 ±<br>0.01389 | 0.2049 ±<br>0.01361 | 0.2013 ±<br>0.01554 | 0.2081 ±<br>0.01637 | 0.1996 ±<br>0.009656 |

Next, visual analysis of the 100th ns snapshot of the MD simulation revealed key residues potentially contributing to the ligand recognition and binding process (Figure 5). JQ1 was shown to form a crucial hydrogen bond with the N140 residue and hydrophobic interactions with I146, W81, L92, and L94. This crucial hydrogen bond has also been seen in the NPC268484, NPC295021, and NPC313112 but is absent in NPC12461 and NPC103230. Hydrophobic interactions were found in all the NP complexes.

**Figure 5.**
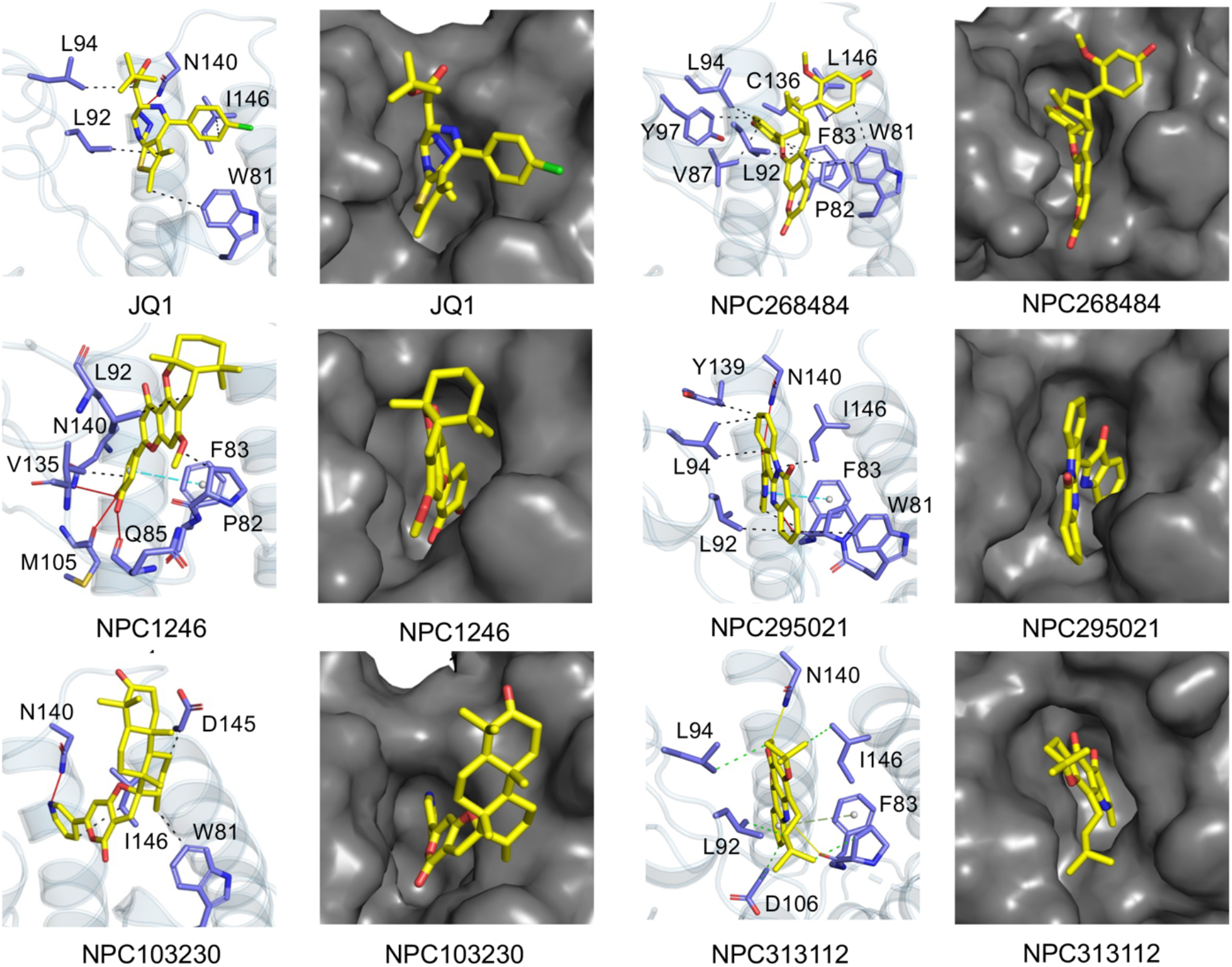
100^th^ ns snapshots from trajectory of all top 5 complexes including JQ1 dotted green line represents hydrophobic interaction and solid yellow line represents hydrogen bonding another dotted dark line represents pi-pi stacking.

### MM/PBSA analysis

In order to calculate a more accurate binding free energy between BRD4-BD1 and NPs, the energy contribution of each residue was calculated by employing the MM/PBSA method. Here, the binding free energy defines the lifetime of the drug molecule inside the binding pocket and the energy contribution by all non-bonded interaction energies between the protein receptor and ligand throughout the 100 ns simulation. Lower binding energy implies better binding and vice versa. The calculated binding free energy between the protein and NPs and its different components like van der Waals forces, electrostatic energy, polar solvation energy, and SASA (solvent accessible surface area) have been listed in Table 4.

**Table 4.** Binding free energy (MM/PBSA) and their components (kJ/mol) of protein complexes

| **Ligands** | **van der Waal**<br>**energy**<br>**kJ/mol** | **Electrostatic**<br>**energy**<br>**kJ/mol** | **Polar solvation**<br>**energy**<br>**kJ/mol** | **SASA energy**<br>**kJ/mol** | **Binding**<br>**energy**<br>**kJ/mol** |
| --- | --- | --- | --- | --- | --- |
| JQ1 | -125.5 ± 22.9 | -8.7 ± 9.6 | 51.5 ± 14.9 | -16.1 ± 2.0 | -98.8 ± 15.9 |
| NCP12461 | -172.0 ± 19.1 | -20.3 ± 9.0 | 118.6 ± 20.7 | -19.0 ± 1.6 | -92.8 ± 14.0 |
| NCP103230 | -122.9 ± 11.9 | -16.9 ± 7.2 | 67.5 ± 12.7 | -15.4 ± 1.1 | -87.8 ± 12.1 |
| NCP268484 | -133.8 ± 30.8 | -19.8 ± 11.8 | 87.1 ± 32.5 | -17.2 ± 3.2 | -96.8 ± 32.1 |
| NCP295021 | -159.6 ± 12.1 | -11.1 ± 6.8 | 104.4 ± 13.0 | -17.7 ± 1.1 | -97.4 ± 11.8 |
| NCP313112 | -170.2 ± 14.0 | -6.3 ± 4.8 | 67.5 ± 9.1 | -18.3 ± 0.8 | -127.3 ± 12.8 |

The binding energy of the BRD4-BD1-JQ1 complex was found to be −96.3 kJ/mol, but some complexes like BRD4-BD1-NCP295021 and BRD4-BD1-NCP313112 have lower binding energies, −131.7 kJ/mol, and −98.4 kJ/mol, respectively (Table 4), and thus, bind more effectively to BRD4-BD1, compared to JQ1. The other three NPs; NCP12461, NCP103230, and NCP268484, show similar or higher binding energies than the reference molecule. This result clearly shows that these 5 NPs has the potential to become better drug molecule than JQ1.

The contribution of each BRD4-BD1 residue to the binding energy in different complexes is depicted in Figure 6. We have selected residues whose energy contribution is lower than −1 kJ/mol. Analysis of the different binding energy plots in Figure 6 reveals that the residues drive recognition and binding in each complex. L92 and I146 contribute the most to the reference molecule JQ1. Other than these two residues, L94, V87, F83, and W81 also contribute significantly to the binding process in JQ1 (Figure 6). The five NPs also show a similar trend to JQ1 regarding the residues facilitating binding. In summary, we selected the top 3 compounds for drug toxicity analysis: candidin (NPC295021), palodesangren B (NPC268484), and buxifoliadin D (NPC313112) (Figure 7).

**Figure 6.**
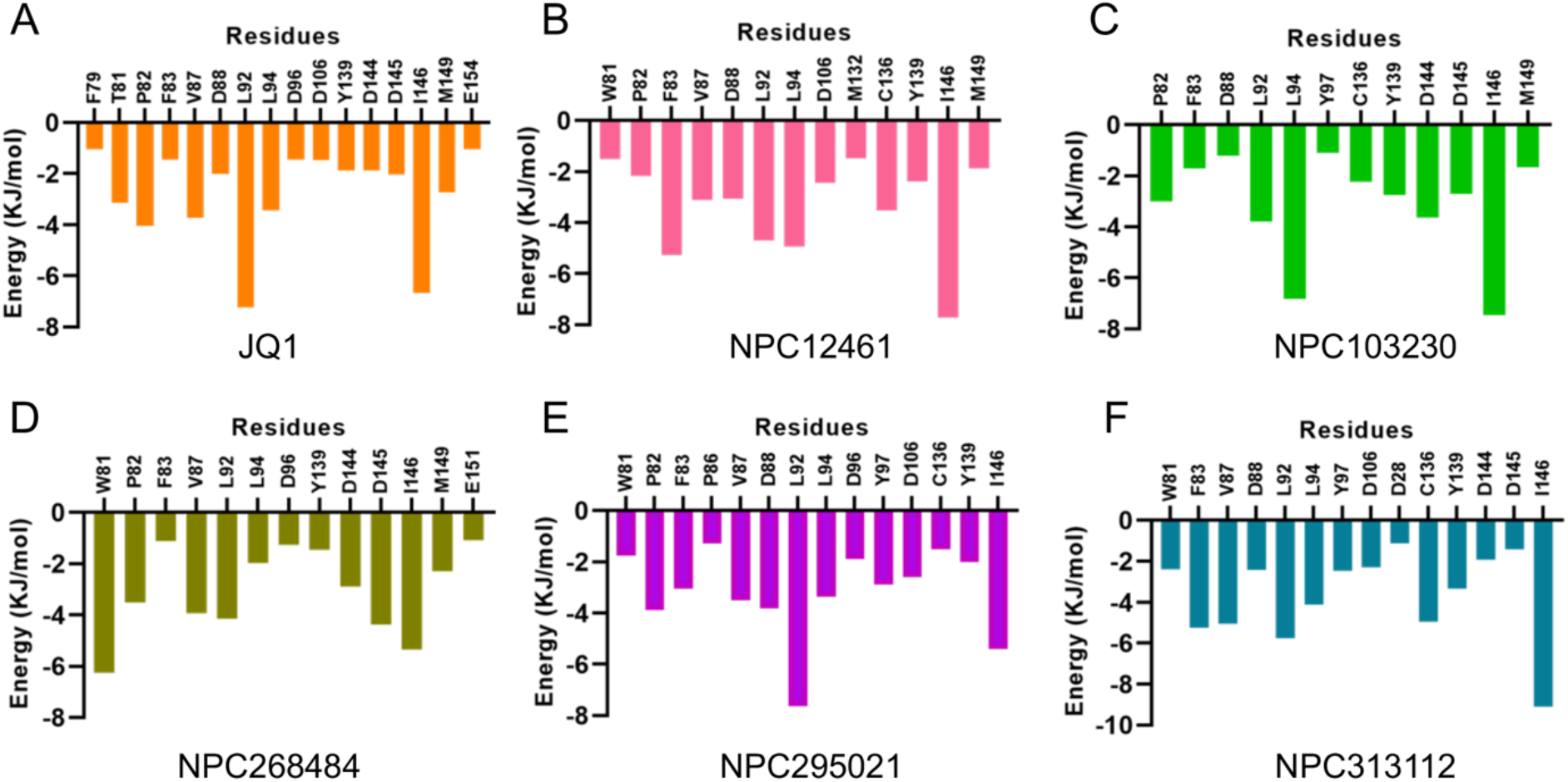
MM/PBSA per residue energy contribution to the binding energy (only residues scoring 1 kJ/mol are shown) for individual compounds.

**Figure 7.**
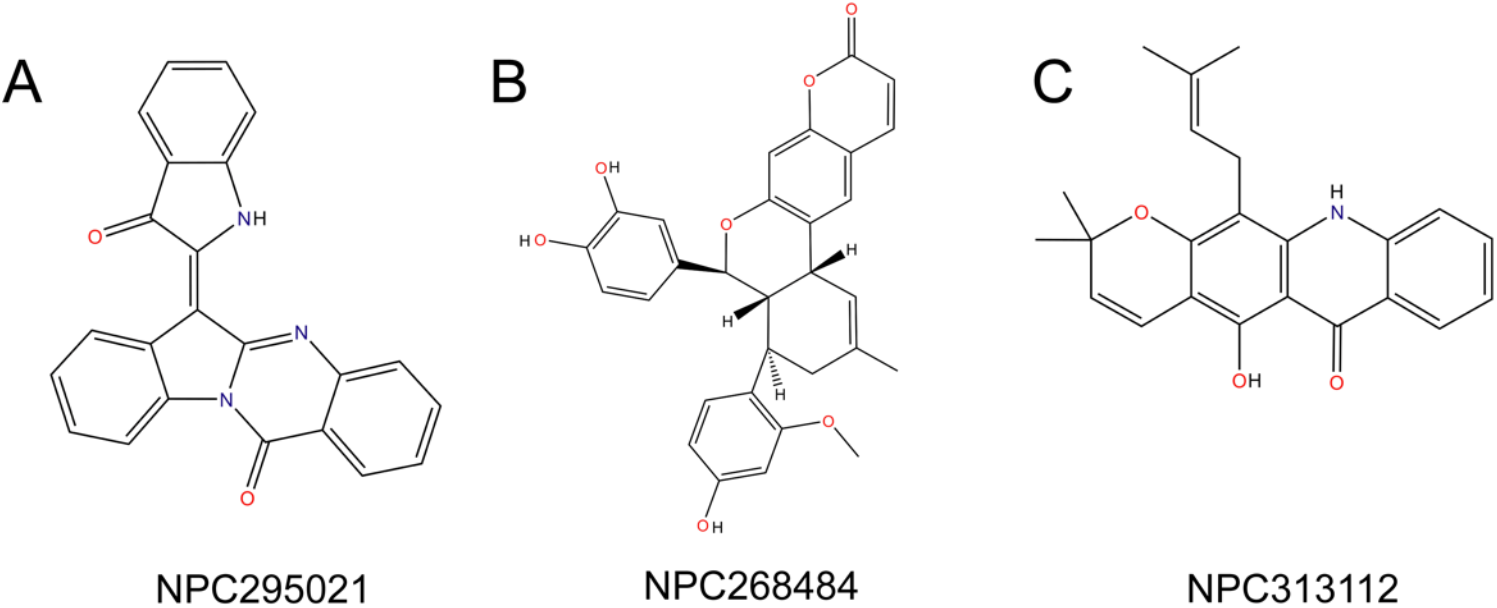
Chemical structures of the top 3 natural products identified in this study.

### Toxicity pattern analysis of top drug candidates

We predicted different toxicity endpoints such as acute toxicity, hepatotoxicity, cytotoxicity, carcinogenicity, mutagenicity, immunotoxicity, adverse effects (Tox21), and toxicity targets were analyzed (Table 5). These results showed that candidin falls into the toxicity class 5 category, while palodesangren B and buxifoliadin D fall into class 4 (the lower the class, the higher the toxicity). The estimated LD50 for candidin, palodesangren B and buxifoliadin D were 300, 500 and 500 mg/kg, respectively. The toxicity radar in Figure 8 illustrates the reliability of positive toxicity results compared to the average of its class. None of the compounds showed adverse effects such as tumorigenic, severe mutagenicity, irritation or reproductive effects. However, candidine showed to be relatively toxic among the four candidates, with little mutagenic and immunotoxic effects (Figure 8 and Table 5).

**Figure 8.**
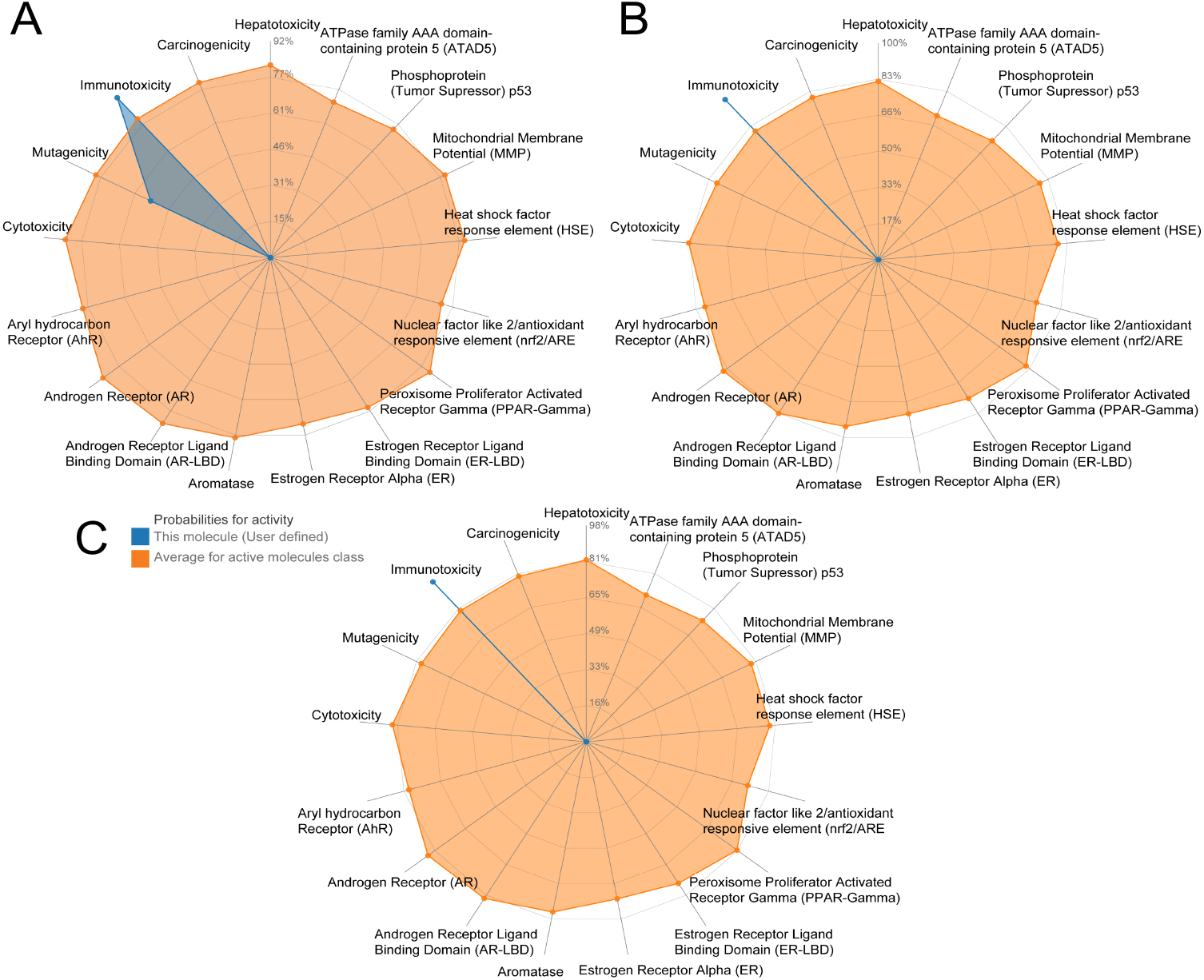
Toxicity patterns of the top three natural products. (A) Candidine, (B) Palodesangren B, (C) Buxifoliadin D.

**Table 5.** Toxicity model reports of the top three candidates

| Classification | Target | Prediction and Probability |  |  |
| --- | --- | --- | --- | --- |
|  |  | NPC29502<br>(Candidine) | NPC268484<br>(Palodesangren B) | NPC313112<br>(Buxifoliadin D) |
| Organ toxicity | Hepatotoxicity | Inactive (0.54) | Inactive (0.66) | Inactive (0.65) |
| Toxicity end points | Carcinogenicity | Inactive (0.5) | Inactive (0.51) | Inactive (0.63) |
| Toxicity end points | Immunotoxicity | Active (0.92) | Active (0.99) | Active (0.97) |
| Toxicity end points | Mutagenicity | Active (0.54) | Inactive (0.61) | Inactive (0.5) |
| Toxicity end points | Cytotoxicity | Inactive (0.68) | Inactive (0.61) | Inactive (0.5) |
| Tox21-Nuclear receptor signalling pathways | Aryl hydrocarbon Receptor (AhR) | Inactive (0.62) | Inactive (0.98) | Inactive (0.7) |
| Tox21-Nuclear receptor signalling pathways | Androgen Receptor (AR) | Inactive (0.85) | Inactive (0.98) | Inactive (0.96) |
| Tox21-Nuclear receptor signalling pathways | Androgen Receptor Ligand Binding Domain (AR-LBD) | Inactive (0.97) | Inactive (0.92) | Inactive (0.97) |
| Tox21-Nuclear receptor signalling pathways | Aromatase | Inactive (0.87) | Inactive (0.83) | Inactive (0.81) |
| Tox21-Nuclear receptor signalling pathways | Estrogen Receptor Alpha (ER) | Inactive (0.87) | Inactive (0.78) | Inactive (0.83) |
| Tox21-Nuclear receptor signalling pathways | Estrogen Receptor Ligand Binding Domain (ER-LBD) | Inactive (0.93) | Inactive (0.86) | Inactive (0.9) |
| Tox21-Nuclear receptor signalling pathways | Peroxisome Proliferator Activated Receptor Gamma (PPAR-Gamma) | Inactive (0.94) | Inactive (0.89) | Inactive (0.9) |
| Tox21-Stress response pathways | Nuclear factor (erythroid-derived 2)-like 2/antioxidant responsive element (nrf2/ARE) | Inactive (0.94) | Inactive (0.8) | Inactive (0.87) |
| Tox21-Stress response pathways | Heat shock factor response element (HSE) | Inactive (0.72) | Inactive (0.8) | Inactive (0.87) |
| Tox21-Stress response pathways | Mitochondrial Membrane Potential (MMP) | Inactive (0.84) | Inactive (0.53) | Inactive (0.55) |
| Tox21-Stress response pathways | Phosphoprotein (Tumor Suppressor) p53 | Inactive (0.68) | Inactive (0.68) | Inactive (0.72) |
| Tox21-Stress response pathways | ATPase family AAA domain-containing protein 5 (ATAD5) | Inactive (0.84) | Inactive (0.75) | Inactive (0.95) |

## Discussion

The past few decades have seen an increase in studies focusing on the role of epigenetic mechanisms in transcriptional regulation and disease. Due to epigenetic mechanisms’ reversibility, epigenetic interventions against diseases such as cancer have started to gain much interest. It is no surprise that researchers are trying to develop drugs that can inhibit the activity of epigenetic proteins to revert and improve the prognosis of diseases without permanently damaging or affecting DNA.

BRD4 is a critical member of the BET family, which has been widely studied due to its well-documented role in several solid and hematopoietic cancers.^*12*, *13*^ Currently, three drug molecules are reported against BRD4-BD1: JQ1, I-BET762, and I-BET151.^*16*, *17*^ These molecules are either not very specific to the BRD4-BD1 protein or have toxic effects on cells. Studies on JQ1 have shown that it causes unwanted damage to normal cells. Particularly in mesenchymal stem cells and their neuronal derivatives, JQ1 inhibited cell proliferation and caused cell death by triggering intrinsic apoptotic pathway thereby effecting their self-renewal which is important for repair and regeneration of damages cells and tissues.^*24*, *49*^ It has also been reported that JQ1 treatment might impair adaptive immune response by suppressing production of IFN-γ in NK cells and CD4+ memory T cells, which is necessary for adaptive immune response against intracellular pathogens.^*50*^ Therefore, it is necessary to identify new, safe and specific inhibitor molecules against BRD4-BD1.

Most inhibitors studies against BRD4 have adopted libraries of synthetic drug molecules. While synthetic products are popular due to low production cost, time effectiveness, stringent regulation, easy quality control, and quick effects, their safety and efficacy have always remained questionable.^*51*^ Natural products, on the other hand, are a good source of safe and effective bioactive molecules that can act as alternatives to synthetic drug candidates. So, for the first time, we decided to work with natural products and reported a new class of BRD4-NP inhibitors, characterized the compounds, and studied the binding dynamics of these novel compounds as a starting point for future natural products inhibitors against BRD4.

In this study, we have employed molecular docking, similarity clustering, pharmacokinetic calculations, MD simulation, and MM/PBSA to find the best NP inhibitors against BRD4-BD1. Using AutoDock vina, we have conducted virtual screening of compounds in the NPASS database targeting the acetyllysine binding site of BRD4-BD1. We identified 10 compounds with binding affinities better than JQ1. Furthermore, the multidimensional clustering and physicochemical characterization gave us the top 5 compounds with drug-like properties. After that, molecular dynamics simulation and MM/PBSA binding free energy calculations revealed that the NPC268484, NPC295021, and NPC313112 compounds are the most stable as compared to the other compounds (Figure 7). Furthermore, best binding inhibitors in the acetyllysine binding cavity are also stable throughout the simulation as they pose a crucial interaction with Asn140 (Figure 5). The structures of all these top NPs are completely different from reported BRD4 inhibitors’ structures. They are NPC268484 (Palodesangren B), NPC295021 (Candidine), and NPC313112 (Buxifoliadine-D), naturally sourced from *Brosimum rubescens* (NPC268484), *Phaius mishmensis, Isatis indigotica, Persicaria tinctorial, Baphicacanthus cusia, Peucedanum praeruptorum* (NPC295021) and *Severinia buxifolia, Atalantia buxifolia* (NPC313112). Our study also revealed the immunotoxic nature of all compounds and having mild mutagenic nature of candidine through computational investigations. Analyzing the similarity distribution with clinically approved drugs from the NPASS server, we found that these NPs belong to a new category of drug molecules (Figure S4 and Table S1). Furthermore, recent reports suggest that candidate Candidine (NPC295021) is a promising drug molecule against SARS-CoV-2 spike protein. Future studies are needed to determine the applicability of these compounds.^*52*^

## Conclusion

Since the discovery of bromodomains in 1992, BRD4 has emerged as one of the most studied bromodomain due to its cancer-related connotations. Multiple attempts have been made to make successful drug candidates. Though some are currently in several stages of clinical trials, they still leave much to be desired, specificity or toxicity. Our high throughput virtual screening (HTVS) study has elucidated 3 top hits: Palodesangren B, Candidine, and Buxifoliadine-D, from the NPASS database. These candidates are completely novel as they do not share any structural or shape similarity with existing inhibitors against BRD4-BD1 and have better binding affinity and stability than them. The results of this study are valuable for the future rational design of better BRD4-BD1 inhibitors from natural sources for various therapies. As explored in this study, these NPs need to be included in drug research targeting BRD4-BD1.

## Acknowledgments

The authors are thankful for the research funding from IISER Kolkata, infrastructural facilities supported by IISER Kolkata. This work was supported by grants from SERB (SRG/2019/000765 and EEQ/2020/000149) and a DBT Ramalingaswami Fellowship (BT/RLF/Re-entry/56/2018) to B.S.

## Statement and Declaration

The authors declare no competing financial interest.

