## Supporting Information for "Identification of novel natural product inhibitors of BRD4 using high throughput virtual screening and MD simulation"

---

**Table S1.** Predicted drug targets for Candidine, Palodesangren B, and Buxifoladin D

| | Target | Common name | Uniprot ID | ChEMBL ID | Target Class | Probability* |
|---|---|---|---|---|---|---|
| NPC29502 (Candidine) | Tyrosine-protein kinase TYK2 | TYK2 | P29597 | CHEMBL3553 | Kinase | 0 |
| | Tyrosine-protein kinase SYK | SYK | P43405 | CHEMBL2599 | Kinase | 0 |
| | Tyrosine-protein kinase LCK | LCK | P06239 | CHEMBL258 | Kinase | 0 |
| | Tyrosine-protein kinase JAK2 | JAK2 | O60674 | CHEMBL2971 | Kinase | 0 |
| | Translocator protein (by homology) | TSPO | P30536 | CHEMBL5742 | Membrane receptor | 0 |
| | Thromboxane-A synthase | TBXAS1 | P24557 | CHEMBL1835 | Cytochrome P450 | 0 |
| | TGF-beta receptor type I | TGFBR1 | P36897 | CHEMBL4439 | Kinase | 0 |
| | Telomerase reverse transcriptase | TERT | O14746 | CHEMBL2916 | Enzyme | 0 |
| | Tankyrase-1 | TNKS | O95271 | CHEMBL6164 | Enzyme | 0 |
| NPC268484 (Palodesangren B) | 5-lipoxygenase activating protein | ALOX5AP | P20292 | CHEMBL4550 | Other cytosolic protein | 0.125687253 |
| | Serine/threonine-protein kinase Aurora- B | AURKB | Q96GD4 | CHEMBL2185 | Kinase | 0.125687253 |
| | Ribosomal protein S6 kinase 1 | RPS6KB1 | P23443 | CHEMBL4501 | Kinase | 0.125687253 |
| | Serine/threonine-protein kinase Aurora- A | AURKA | O14965 | CHEMBL4722 | Kinase | 0.125687253 |
| | Estradiol 17-beta-dehydrogenase 2 | HSD17B2 | P37059 | CHEMBL2789 | Enzyme | 0.125687253 |
| | Estradiol 17-beta-dehydrogenase 1 | HSD17B1 | P14061 | CHEMBL3181 | Enzyme | 0.125687253 |
| | Gastrin releasing peptide receptor | GRPR | P30550 | CHEMBL4959 | Family A G protein-coupled receptor | 0.125687253 |
| NPC313112 (Buxifoliadin D) | 11-beta-hydroxysteroid dehydrogenase 1 | HSD11B1 | P28845 | CHEMBL4235 | Enzyme | 0.113285953 |
| | 3-phosphoinositide dependent protein kinase-1 | PDPK1 | O15530 | CHEMBL2534 | Kinase | 0.113285953 |
| | 5-lipoxygenase activating protein | ALOX5AP | P20292 | CHEMBL4550 | Other cytosolic protein | 0.113285953 |
| | ADAMTS5 | ADAMTS5 | Q9UNA0 | CHEMBL2285 | Protease | 0.113285953 |
| | Adenosine A1 receptor | ADORA1 | P30542 | CHEMBL226 | Family A G protein -coupled receptor | 0.113285953 |
| | Adenosine A2a receptor | ADORA2A | P29274 | CHEMBL251 | Family A G protein -coupled receptor | 0.113285953 |
| | Adenosine A2b receptor | ADORA2B | P29275 | CHEMBL255 | Family A G protein -coupled receptor | 0.113285953 |
| | Alkaline phosphatase, tissue-nonspecific isozyme | ALPL | P05186 | CHEMBL5979 | Enzyme | 0.113285953 |
| | Alpha-1a adrenergic receptor | ADRA1A | P35348 | CHEMBL229 | Family A G protein-coupled receptor | 0.113285953 |

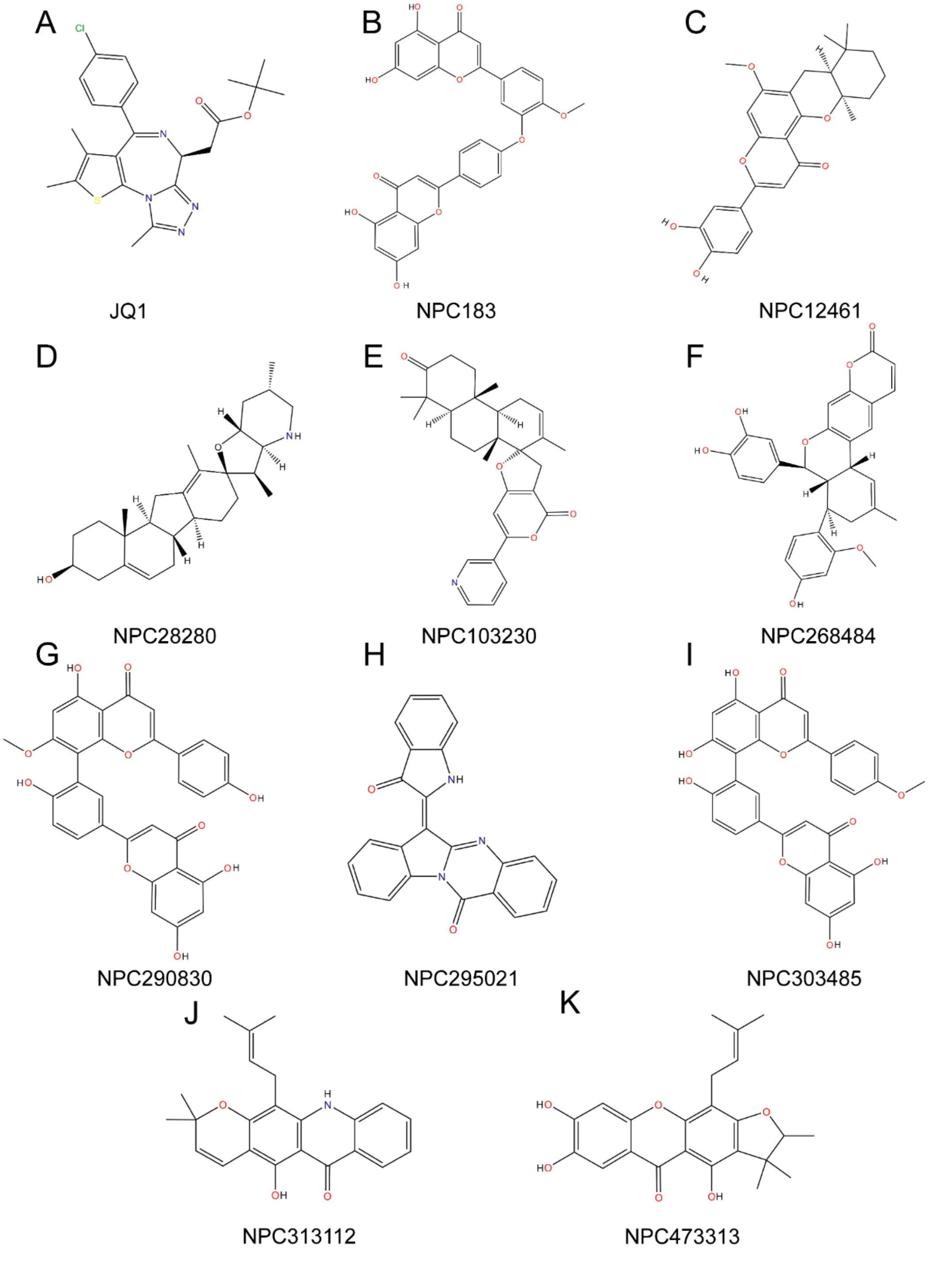


**Figure S1.** Chemical structures and NPASS Ids of the potential ligands in the Kac binding site.

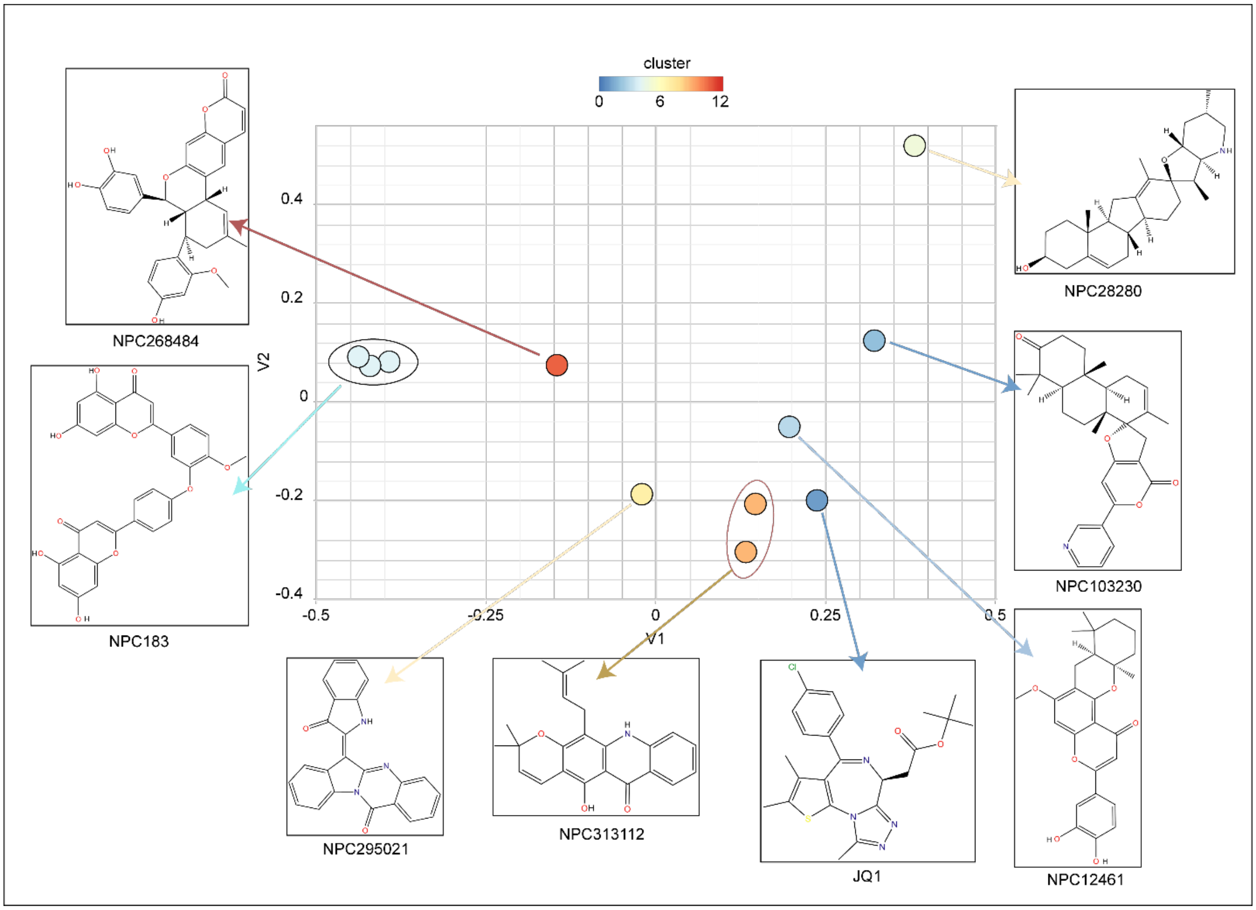


**Figure S2**. Clustering of all the compounds Multidimensional Scaling (MDS) and from each cluster one example structure is showing different colour coat represents different chemical group Tanimoto coefficient <0.4 and same colour coat represents > 0.4.

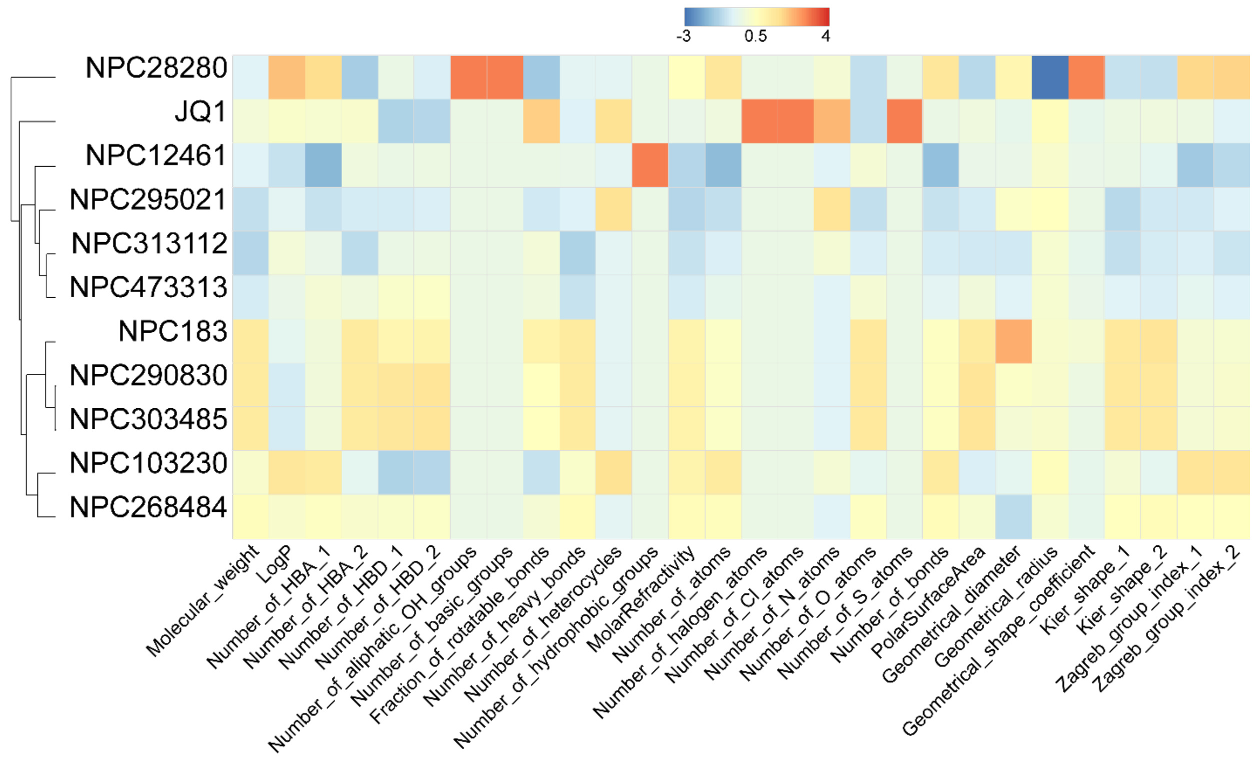


**Figure S3**. Hierarchical clustering Joe lib descriptor heatmap of top 10 compounds.

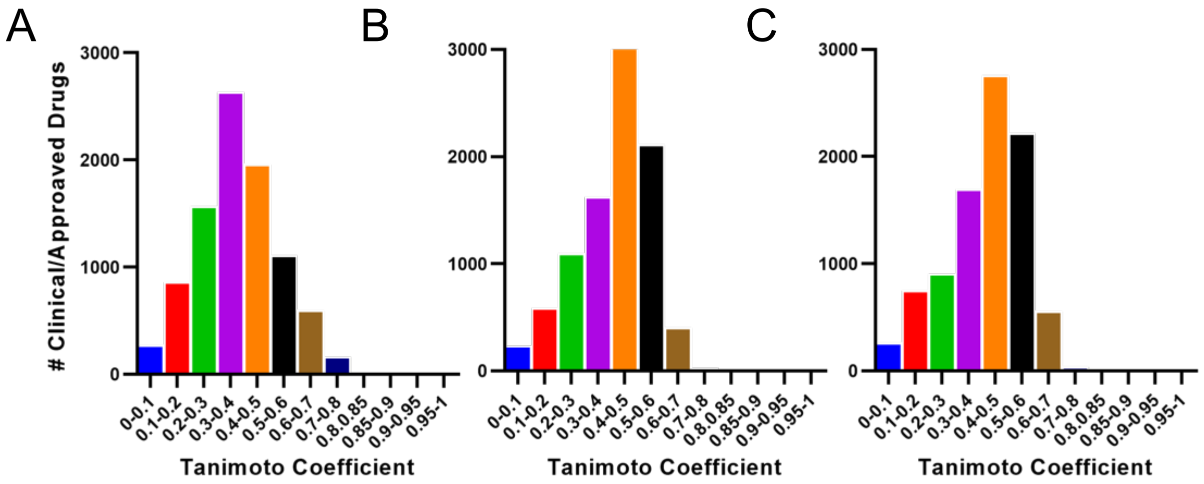


**Figure S4.** Similarity distribution of top 3 compounds with clinically approved drugs and all the molecules identified are unique.
